# High-resolution cryo-EM structures of small protein–ligand complexes near the theoretical size limit

**DOI:** 10.1101/2025.06.30.662489

**Authors:** Kunwoong Park, Youngki Yoo, Hyunbum Jeon, Kiju Choi, Eunju Kwon, Hyun-Ho Lim, Dong Young Kim, Kyoung Tai No

## Abstract

Cryo-electron microscopy (cryo-EM) is widely used to determine macromolecular structures at atomic resolution. The theoretical size lower limit of particles for cryo-EM analysis is 38 kDa, limited by factors such as contrast and particle alignment accuracy. To date, no cryo-EM structures have been reported for proteins near this size limit. This study presents cryo-EM structures of two protein-ligand complexes around 40 kDa. The structure of maltose-binding protein (43 kDa) was determined at 2.32 Å resolution, clearly revealing the bound maltose and water molecules. Additionally, the kinase domain of human PLK1 (37 kDa), slightly below the theoretical limit, was determined at 3.04 Å resolution, allowing the identification of the bound ligand, onvansertib. These findings demonstrate that cryo-EM can be effectively employed for structure determination and structure-based drug screening of small proteins or domains.

## INTRODUCTION

Cryo-EM is an advanced technique used to determine the structures of biological macromolecules at atomic resolution. It has become a primary tool in structural biology because it enables structural analysis of frozen macromolecules without requiring crystallization, a major barrier in protein crystallography. Traditionally, achieving atomic-resolution with cryo-EM has been limited by particle size. However, recent advancements in detector sensitivity, motion correction, and data processing algorithms have made significant improvements^1^. The theoretical minimum particle size for single-particle reconstruction is estimated to be 38 kDa, based on factors such as radiation damage and the signal-to-noise ratio^2^. Recently, two cryo-EM structures of RNA particles near the size limit have been successfully determined at atomic resolution. Structure of the SAM-IV riboswitch RNA (40 kDa) was determined at 3.7 Å resolution using conventional cryo-EM single particle reconstruction, and structure of the ACA2 dimer complexed with RNA (40 kDa) was determined at 2.6 Å resolution using a deep-learning based single particle reconstruction method^3-5^. RNA molecules contain a phosphate group with strong electron scattering, providing improved contrast for visualization compared to proteins^6^. This characteristic makes it advantageous for high-resolution structural analysis of RNA. The smallest protein analyzed by cryo-EM to date is NADPH oxidase, with a molecular weight of 46 kDa^7^. In this study, we present cryo-EM structures of two small protein-ligand complexes. The maltose-binding protein (MBP, 43 kDa) complexed with maltose was determined at a resolution of 2.32 Å, and the human polo-like kinase 1 kinase domain (hPLK1_KD_, 37 kDa) complexed with the anti-cancer compound onvansertib was determined at a resolution of 3.04 Å. Both structures were determined using a conventional single-particle reconstruction method without symmetry enforcement. These structural analyses demonstrate the potential of conventional cryo-EM methods to determine high-resolution structures and facilitate structure-based drug discovery for small protein-ligand complexes, extending beyond the theoretical size limit.

## RESULTS AND DISCUSSION

### Cryo-EM structure of MBP complexed with maltose

MBP is commonly used as a fusion tag for the expression of recombinant proteins, as it enhances the solubility of target proteins and simplifies purification via maltose-binding affinity chromatography^8,9^. In this study, we purified MBP and determined its cryo-EM structure using conventional single-particle reconstruction methods. Recombinant MBP was expressed in *Escherichia coli* using an MBP-fusion vector system and purified through maltose-binding affinity chromatography followed by size-exclusion chromatography (SEC). Cryo-EM micrographs of purified MBP exhibited high particle contrast even at low defocus values (-0.4 to -1.2 μm), with well-defined high-frequency Thon-rings and low astigmatism (mean value: 181.5 ± 42 Å), indicative of thin, uniform ice and high-quality data suitable for high-resolution reconstruction (**Fig. S1**). A total of 2,652 micrographs were collected using a 300 kV cryo-transmission electron microscope, from which 6,021,153 particles were extracted. The cryo-EM map of MBP was reconstructed at an overall resolution of 2.32 Å using 517,863 particles (**Fig. S2**). The atomic model built from the cryo-EM map showed no outliers in the Ramachandran plot (**Table S1**). The final structure includes one MBP monomer, one maltose molecule, and 492 water molecules. The cryo-EM map accurately identified all residues and ligands in the atomic model (**Figs. 1A–1C**).

**Figure 1.**
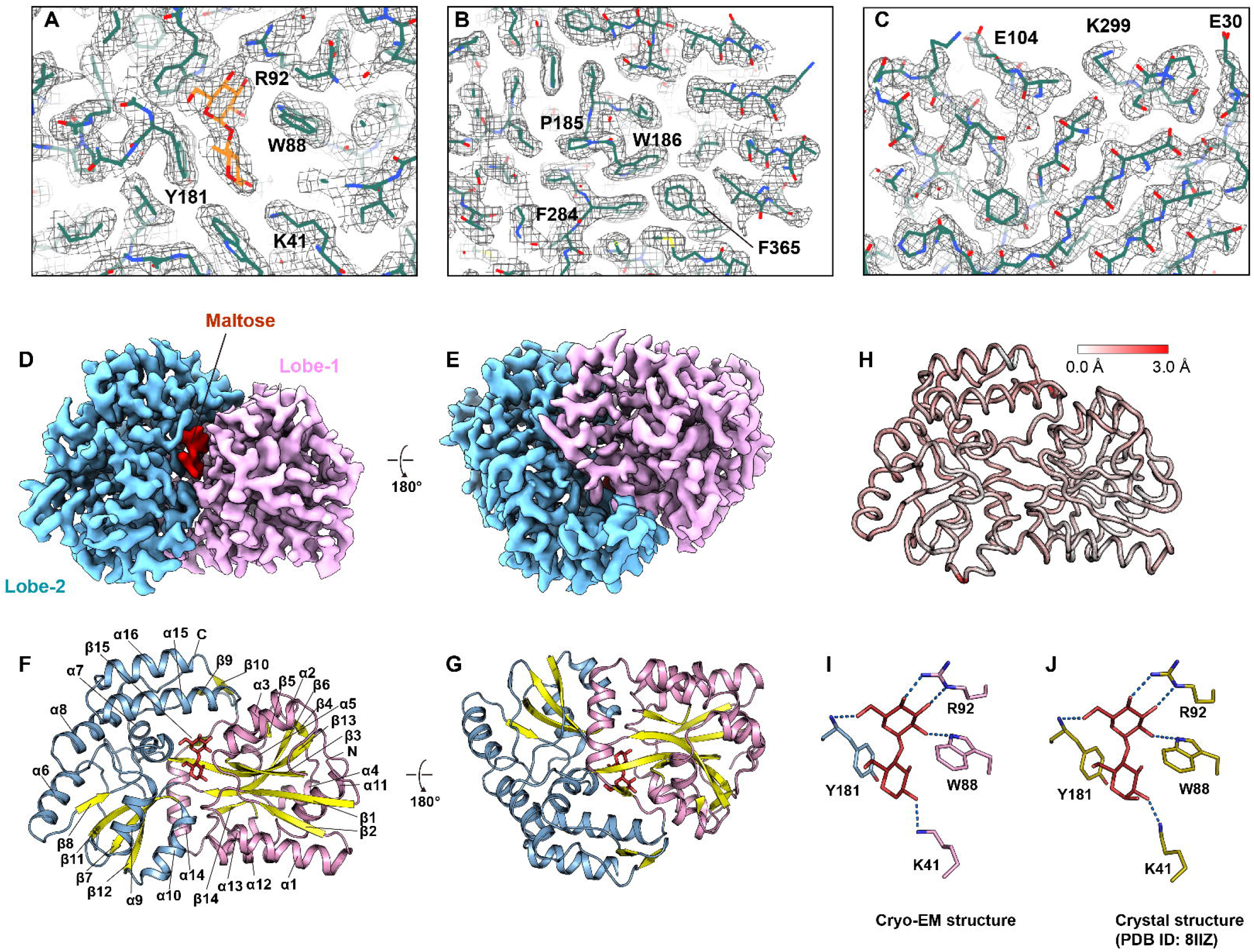
Cryo-EM structure of the MBP–maltose complex. (A–C) Close-up views of the cryo-EM density map and its atomic model at the maltose binding site (A), the crevice between Lobe-1 and Lobe-2 (B), and the surface of Lobe-1 (C). (D, E) Cryo-EM maps of MBP shown in two different orientations. Lobe-1, Lobe-2, and maltose are colored pink, blue, and red, respectively. (F, G) Cartoon models depicted in the same orientations as (D) and (E). Secondary structures are labeled α1–α16 for helices and β1–β15 for strands. (H) Cα-trace model of MBP showing structural differences between the cryo-EM and crystal structures. A white-to-red color spectrum represents root-mean-square (RMS) values ranging from 0.0 Å to 3.0 Å. (I, J) Stick models illustrating interactions between MBP and maltose. (I) Cryo-EM structure. (J) Crystal structure of MBP (PDB ID: 8IIZ).

In the cryo-EM structure, MBP exhibits a two-lobe conformation (**Figs. 1D and 1E**). Lobe-1 features an α/β fold characterized by a central β-sheet with strands arranged in the order β2, β1, β3, β13, β6, and β5/β14, with peripheral helices (α1**–**α5 and α11**–**α14). This lobe also includes β4 and β15 strands that interact with β13. Similarly, Lobe-2 forms an α/β fold with a central β-sheet composed of β8, β11, β7, and β12, along with peripheral helices (α6–α10 and α15–α16) and an additional two-stranded β-sheet formed by β9 and β10 (**Figs. 1F and 1G**). The cryo-EM structure of MBP closely resembles its crystal structure (PDB ID: 8IIZ)^10,11^ **(Fig. 1H)**. Superimposing both structures yields a root-mean-square deviation (RMSD) of 0.6 Å for 367 Cα atoms. In the cryo-EM structure, a maltose molecule is bound within the cleft between the two lobes, forming hydrogen bonds with residues K41, W88, R92, and Y181 (**Fig. 1I**). These interactions are consistent with those observed in the crystal structure of MBP (**Fig. 1J**). This analysis suggests that the cryo-EM structure can elucidate structural features of the MBP–maltose complex with accuracy comparable to its high-resolution crystal structure.

### Cryo-EM Structure of hPLK1_KD_ complexed with onvansertib

PLK1 is a serine/threonine protein kinase that activates the G2/M transition and regulates meiosis during the cell cycle^12^. As a proto-oncogene, PLK1 is frequently overexpressed in tumor cells, making it a key target for cancer drug development^13^. Onvansertib is a selective PLK1 inhibitor that competitively binds to the ATP binding site of the kinase^14^. To investigate the potential of cryo-EM for analyzing small protein–ligand complexes, we conducted a cryo-EM structural analysis of the hPLK1_KD_ (residues 13–345) in complex with onvansertib, a complex with an approximate size of 37 kDa. hPLK1_KD_ was expressed using a baculovirus system and purified through immobilized metal affinity chromatography followed by SEC. The purified protein was mixed with onvansertib and applied to cryo-EM grids. Similar to MBP samples, the cryo-EM micrographs of hPLK1_KD_ exhibited high particle contrast, well-defined Thon rings, and low astigmatism (mean value: 140 ± 78 Å), supporting the quality of the dataset for high-resolution reconstruction (**Fig. S3**). A total of 4,053 micrographs were collected using a 300 kV cryo-transmission electron microscope. From these, the cryo-EM map of the hPLK1_KD_–onvansertib complex was reconstructed at an overall resolution of 3.04 Å from 134,422 particles, without imposing any symmetry (**Fig. S4**). Angular distribution plot analysis revealed that particle orientation was biased toward a single direction, potentially limiting the overall resolution and resulting in weak density in certain regions (**Fig. S4C**). Nevertheless, the map quality was sufficient to build an atomic model of hPLK1_KD_ and to fit the onvansertib ligand (**Figs. 2A–2C**).

**Figure 2.**
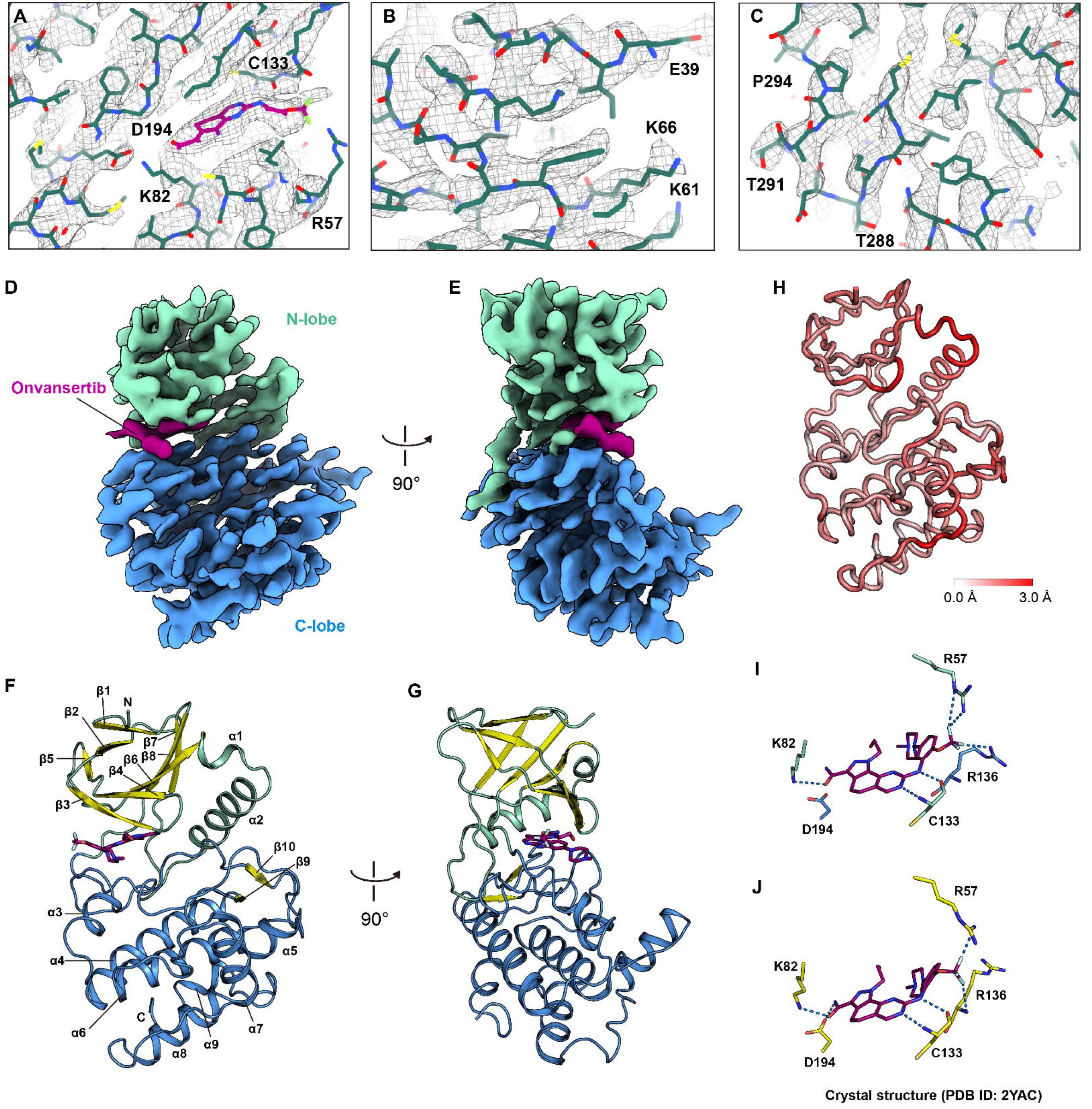
Cryo-EM structure of the hPLK1_KD_–onvansertib complex. (A–C) Close-up views of the cryo-EM density map and its atomic model at the onvansertib binding site (A), the N-Lobe surface (B), and the C-Lobe surface (C). (D, E) Cryo-EM maps of hPLK1_KD_ shown in two different orientations. The N-lobe, C-lobe, and onvansertib are colored cyan, blue, and magenta, respectively. (F, G) Cartoon models depicted in the same orientations as (D) and (E). Secondary structures are labeled α1–α9 for helices and β1–β10 for strands. (H) Cα-trace model of hPLK1_KD_ showing structural differences between the cryo-EM and crystal structures. A white-to-red color spectrum represents root-mean-square (RMS) values ranging from 0.0 Å to 3.0 Å. (I, J) Stick models illustrating interactions between hPLK1_KD_ and onvansertib in the cryo-EM structure (I) and the crystal structure (PDB ID: 2YAC).

In the cryo-EM structure, hPLK1_KD_ exhibits a two-lobe conformation (**Figs. 2D and 2E**). The N-lobe features a β-sandwich composed of two β-sheets interspersed with helices (α1 and α5). One β-sheet is composed of β1, β2, and β5, and the other comprises β3, β4, β6, β7, and β8. The C-lobe forms an α-helical fold with a short two-stranded β-sheet formed by β9 and β10 (**Figs. 2F and 2G**). The cryo-EM structure of hPLK1_KD_ closely resembles its crystal structure (PDB ID: 2YAC)^14^ **(Fig. 2H)**. When the two structures are superimposed, the RMSD for 268 Cα atoms is 1.0 Å. In the cryo-EM structure, onvansertib is bound within the creft between the two lobes, forming hydrogen bonds with residues R57, K82, C133, and R136 of hPLK1_KD_ (**Fig. 2I**). These interacting residues are consistent with those observed in the crystal structure of hPLK1_KD_, except that D194 is slightly distant to form a hydrogen bond with onvansertib (**Fig. 2I and 2J**). This comparison suggests that the cryo-EM structure can elucidate the structural features of the hPLK1_KD_–onvansertib complex as accurately as the high-resolution crystal structure.

### Implications of cryo-EM determination of small protein-ligand complexes

To overcome size limitations in cryo-EM structural analysis, various strategies have been developed, including phase-plate imaging^15,16^, graphene supports^17^, particle symmetry, and engineered scaffolds^18,19^. Although these techniques enhance alignment accuracy and improve the signal-to-noise ratio, they often introduce technical complexity or require extensive structural modification. This study presents high-resolution cryo-EM structures of small protein–ligand complexes of 43 and 37 kDa. The cryo-EM map of the MBP–maltose complex shows a clear density map for the protein, ligand, and coordinated water molecules. Similarly, the map of the hPLK1_KD_–onvansertib complex provides sufficient resolution for model building of the protein and ligand. Notably, this represents the smallest cryo-EM structure reported to date, at 37 kDa, surpassing the previously assumed theoretical size limit. Our findings demonstrate that high-resolution cryo-EM structures of asymmetric proteins as small as 40 kDa can be obtained using conventional cryo-EM techniques, without relying on molecular symmetry, scaffolding, or specialized equipment. These results highlight the potential of conventional cryo-EM as a powerful method for elucidating the structures of small proteins and domains bound to chemical drug candidates. Moreover, they suggest that structural analysis may be extended to even smaller macromolecules than the theoretical minimum size. As image contrast and reconstruction algorithms continue to advance, high-resolution cryo-EM analysis of increasingly small and challenging targets is expected to become more routine.

## METHODS

### Protein expression and purification of MBP

MBP was purified as a byproduct during the purification of a Zebrafish PLK1_KD_. DNA encoding the target protein, which included an N-terminal Tobacco etch virus (TEV) protease cleavage sequence, was inserted between the *NdeI* and *BamHI* restriction sites of the pMal-c5x plasmid (Addgene, Watertown, MA, USA). The plasmid was introduced into *E. coli* BL21 (DE3) cells (Thermo Fisher Scientific, Waltham, MA, USA). The transformed cells were cultured in 1 L of Luria-Bertani (LB) medium (Gibco, Waltham, MA, USA) at 37 °C. When the optical density at 600 nm reached 0.8, the culture was transferred to a cold room and cooled for 1 h. Protein expression was then induced by adding 1 mM isopropyl β-D-thiogalactopyranoside. After culturing at 16°C for 18 h, the cells were harvested by centrifugation at 3,000 × *g* for 10 min, resuspended in buffer A (50 mM Tris-HCl pH 7.5, 400 mM NaCl, and 3 mM DTT), and lysed by sonication. The cell lysate was loaded onto Dextrin-Sepharose resin (Cytiva, Marlborough, MA, USA) and eluted with buffer B (50 mM Tris-HCl pH 7.5, 400 mM NaCl, 3 mM DTT, and 10 mM maltose). The eluate was treated with TEV protease and incubated overnight at 4°C to remove C-terminal flexible residues. TEV protease was then removed by passing the solution through Ni-NTA resin (Qiagen, Venlo, Limburg, Netherlands). MBP was further purified by size-exclusion chromatography using a Superdex-200 increase 10/300 GL column (Cytiva) equilibrated with buffer C (50 mM Tris-Cl pH 7.5 and 400 mM NaCl). The purified MBP was concentrated to a final concentration of 1 mg/mL for cryo-EM analysis.

### Protein expression and purification of hPLK1_KD_

DNA encoding hPLK1_KD_ (residues 13–345, containing the T210V mutation) was cloned into the pFastBac1 vector along with a sequence for an N-terminal 6×His tag followed by a TEV protease cleavage site. Recombinant baculovirus was generated using the Bac-to-Bac system (Invitrogen, Carlsbad, CA, USA) and used to infect Sf9 insect cells. The cells were cultured in Sf-900 SFM medium (Gibco) at 27°C for 72 h. Following incubation, cells were harvested by centrifugation at 3,000 × *g* for 10 min, resuspended in buffer D (50 mM HEPES pH 7.5, 500 mM NaCl, 20 mM imidazole, 1 mM β-mercaptoethanol, 1% NP-40, and 1 mM PMSF), and lysed by sonication. The cell lysate was centrifuged at 20,000 × *g* for 30 min, and the supernatant was loaded onto Ni-NTA resin (Qiagen). The resin was washed with buffer E (50 mM HEPES pH 7.5, 500 mM NaCl, 20 mM imidazole, and 1 mM β-mercaptoethanol), and the bound protein was eluted using buffer F (50 mM HEPES pH 7.5, 500 mM NaCl, 300 mM imidazole, and 1 mM β-mercaptoethanol). The eluted protein was incubated with TEV protease overnight at 4°C to remove the 6×His tag. Finally, hPLK1_KD_ was purified through SEC using a HiLoad 26/600 Superdex 200 pg column (Cytiva) equilibrated with buffer G (50 mM HEPES pH 7.5, 150 mM NaCl, and 5 mM TCEP).

### Cryo-EM data collection

UltrAuFoil R1.2/1.3 300-mesh gold grids (SPI Supplies, West Chester, PA, USA) were subjected to glow discharge at 15 mA for 1 min. Following this treatment, 3 μL of MBP at 1 mg/mL was applied to the grids. The grids were blotted for 5 s using humidity-saturated Whatman No. 1 filter paper (Cytiva) and plunge-frozen in liquid ethane using a Vitrobot Mark IV (Thermo Fisher Scientific). Micrographs were obtained using a 300 kV Titan Krios microscope (Thermo Fisher Scientific), equipped with a Falcon 4 direct electron detector and a Selectris-X energy filter with a 10-eV slit width. The micrographs were collected at a pixel size of 0.7052 Å, corresponding to a nominal magnification of 165,000×. The total electron dose was 50 e^−^/Å^2^, and the nominal defocus values ranged from −0.4 to −1.8 μm. For the preparation of the grid specimen of the hPLK1_KD_-onvansertib complex, purified hPLK1_KD_ was mixed with onvansertib at a molar ratio of 1:4. Grid preparation and micrograph collection followed the same procedures as for MBP, with the exception that the nominal defocus values ranged from −0.5 to −2.1 μm.

### Data processing and structure determination

Image processing was performed using cryoSPARC (v.4.4.0) software^20^. Raw micrograph movies were motion-corrected using Patch Motion Correction, and defocus values for each micrograph were estimated using Patch CTF Estimation. To reconstruct the cryo-EM map of MBP, particles were identified using the Template Picker with templates generated from a previous pilot data set. A total of 6,021,153 particles were picked from 2,653 micrographs and extracted with a box size of 300 pixels for two-dimensional (2D) classification. After multiple rounds of 2D classification, 551,521 particles were selected. A separate set of particles was re-picked from the micrographs using the TOPAZ program^21^ in the cryoSPARC platform, yielding 395,799 extracted particles. The two particle sets were combined, and poor-quality particles and duplicates were removed through repeated rounds of 2D classification, resulting in a final dataset of 758,094 particles. Three ab initio models were reconstructed from this combined set, and 593,843 particles corresponding to two of the three classes were subjected to heterogeneous, non-uniform, and local refinement. The final map of MBP was reconstructed from 517,853 particles, with an estimated overall resolution of 2.32 Å, determined by the gold-standard Fourier shell correlation (FSC) at the 0.143 criterion.

To reconstruct the cryo-EM map of the hPLK1_KD_–onvansertib complex, a total of 8,352,733 particles were picked from 4,053 micrographs using the Template Picker, with templates obtained from an initial dataset. These particles were extracted with a box size of 350 pixels for 2D classification. After multiple rounds of 2D classification, a refined set of 577,458 particles was selected. Additionally, another set of 2,118,292 particles was extracted from the micrographs using TOPAZ. The two particle sets were merged, and poor-quality particles and duplicates were removed through repeated 2D classification, resulting in a final selection of 1,476,515 particles.

To generate a template for particle classification, ab initio models were reconstructed from the combined set. A total of 8,352,733 initially picked particles were classified through three rounds of heterogeneous refinement using these ab initio models, yielding a clean subset of 435,356 particles. This subset was used for a second ab initio reconstruction. The final map was generated from 134,422 particles after *ab initio* reconstruction, 3D variability analysis, non-uniform refinement, reference-based motion correction, and local refinement, with an estimated overall resolution of 3.04 Å.

### Atomic model building and structure analysis

Atomic models of MBP and hPLK1_KD_ were built by fitting their crystal structures into the reconstructed cryo-EM maps. The crystal structures of MBP (PDB ID: 1FQC) and hPLK1_KD_ were fitted into the cryo-EM maps using ChimeraX^22^, rebuilt by manual tracing in COOT^23^, and refined using Phenix.refinement^24,25^. Water molecules were found using Phenix.douse^25^. Cryo-EM data collection and refinement statistics are summarized in Table S1. Figures were generated using ChimeraX^22^ and PyMOL^26^.

## DATA AVAILABILITY

The final coordinates and cryo-EM maps supporting the findings of this study have been deposited in the Worldwide Protein Data Bank (www.wwpdb.org) under the following accession codes: PDB IDs (xxxx and xxxx) and EMDB IDs (EMD-xxxxx and EMD-xxxxx).

## AUTHOR CONTRIBUTIONS

K.P., Y.Y., H.J., K.C., and H.K. performed the experiments. K.P., E.K., H.L., D.Y.K., and K.T.N. analyzed the data. K.P., D.Y.K., and K.T.N. wrote the manuscript.

