## Supplementary material for "High-resolution cryo-EM structures of small protein–ligand complexes near the theoretical size limit": Supplemantal Figures 1-4 and Table 1

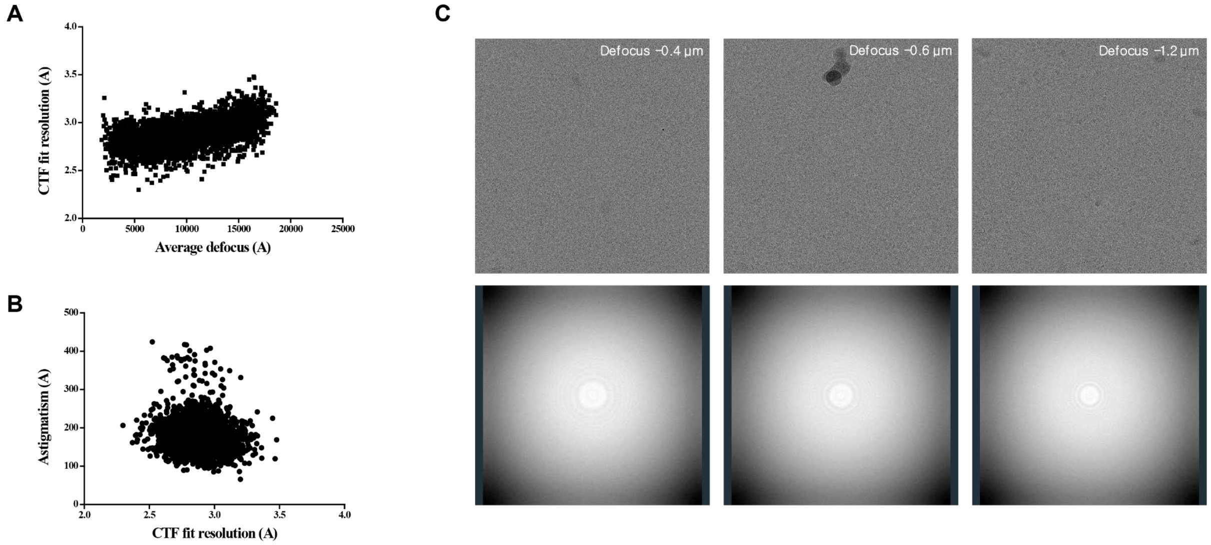


**Figure S1. Evaluation of cryo-EM data quality for the MBP–maltose complex.** (A) Plot showing the correlation between CTF fit resolution and average defocus value. (B) Plot showing the correlation between astigmatism and CTF fit resolution. (C) Representative raw micrographs (upper panels) and corresponding Thon ring patterns (lower panels) collected at defocus values of −0.4 µm, −0.6 µm, and −1.2 µm.

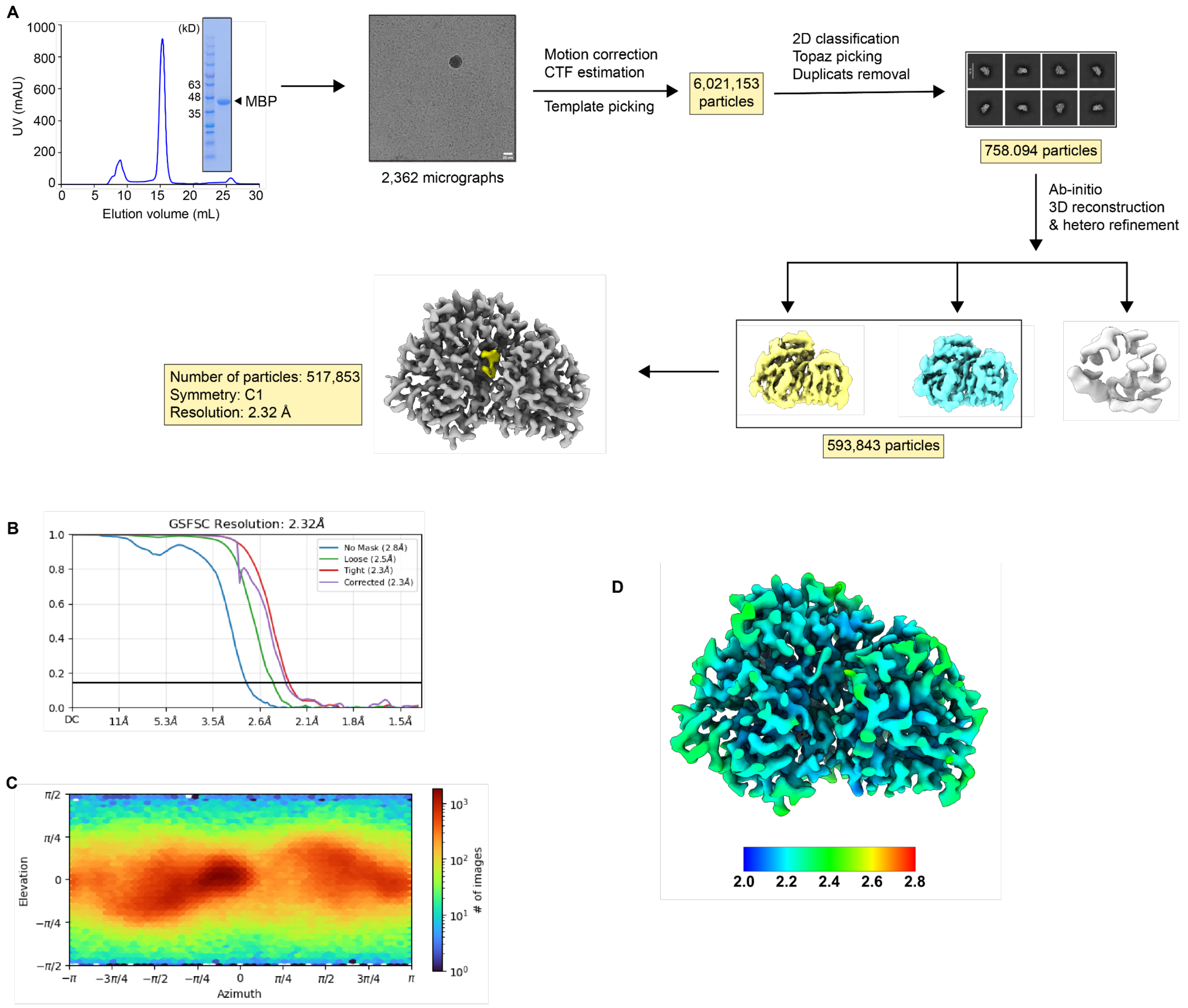


**Figure S2. Cryo-EM data analysis of the MBP–maltose complex.** (A) Workflow for single-particle reconstruction of the MBP–maltose complex. (B) Gold-standard Fourier shell correlation (FSC) curves, with an overall resolution estimated at an FSC value of 0.143. (C) Angular distribution of particle projections. (D) Cryo-EM map showing local resolution.

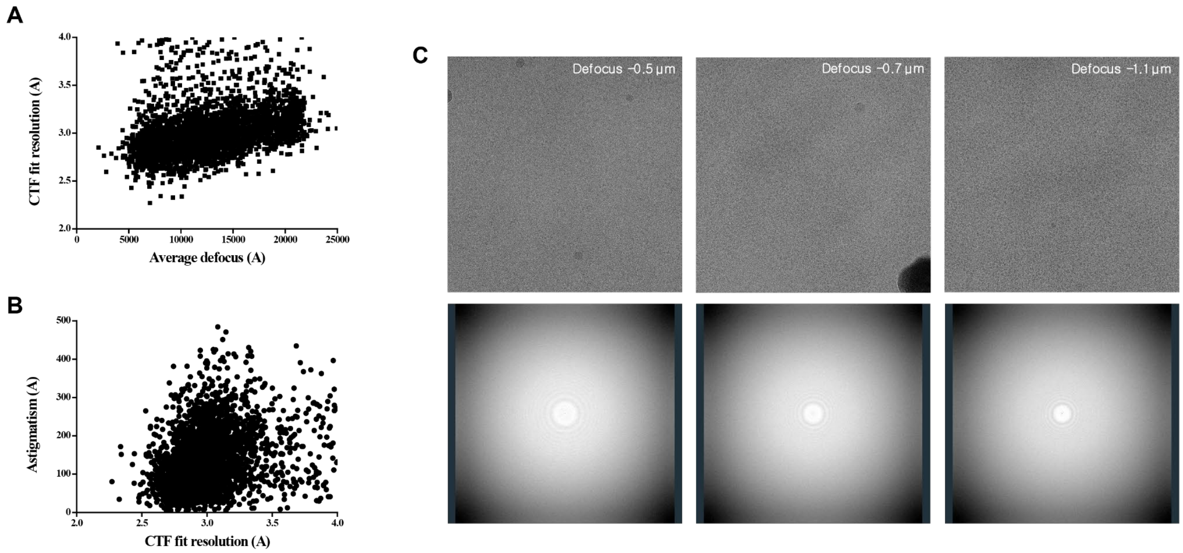


**Figure S3. Evaluation of cryo-EM data quality for the $hPLK1_{KD}$–onvansertib complex.** (A) Plot showing the correlation between CTF fit resolution and average defocus value. (B) Plot showing the correlation between astigmatism and CTF fit resolution. (C) Representative raw micrographs (upper panels) and corresponding Thon ring patterns (lower panels) collected at defocus values of −0.5 μm, −0.7 μm, and −1.1 μm.

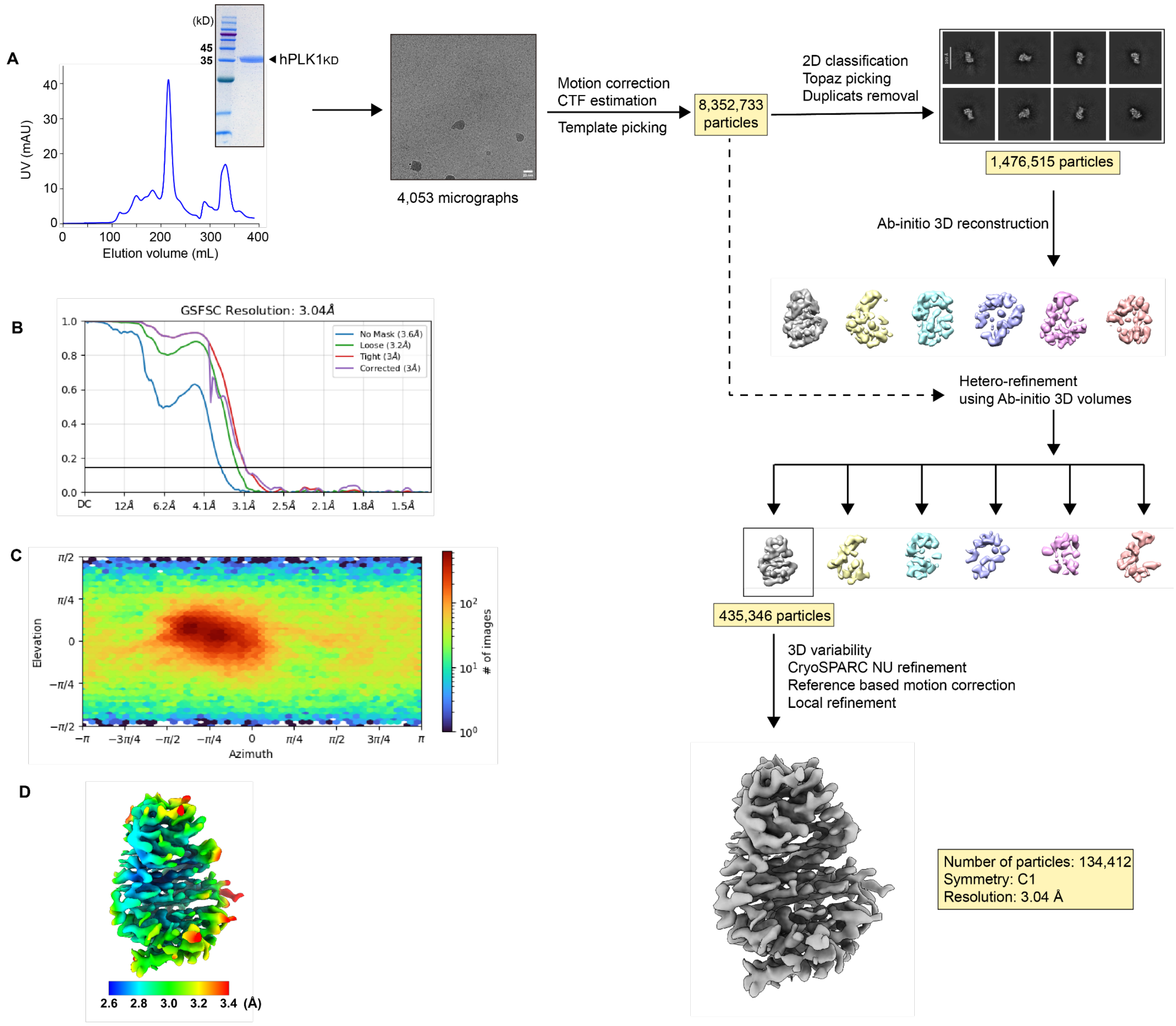


**Figure S4. Cryo-EM data analysis of the hPLK1$_{KD}$–onvansertib complex.** (A) Workflow for single-particle reconstruction of the hPLK1$_{KD}$–onvansertib complex. (B) Gold-standard Fourier shell correlation (FSC) curves, with an overall resolution estimated at an FSC value of 0.143. (C) Angular distribution of particle projections. (D) Cryo-EM map showing local resolution.

**Table S1. Cryo-EM data collection and refinement statistics.**

| **Data set** | MBP-maltose | hPLK1$_{KD}$-onvansertib |
|---|---|---|
| **Data collection** | | |
| Microscope | FEI Titan Krios | FEI Titan Krios |
| Detector | Falcon4 | Falcon4 |
| Magnification | 165,000 | 165,000 |
| Voltage (kV) | 300 | 300 |
| Electron Exposure ($e^-$/Å$^2$) | 60 | 60 |
| Defocus Range (μm) | -0.4 to -1.8 | -0.5 to -2.1 |
| Pixel size (Å) | 0.7052 | 0.7052 |
| No. frames/movie | 60 | 60 |
| Total exposure time (sec) | 4.16 | 4.59 |
| Number of movies | 2,643 | 4.053 |
| **Map reconstruction** | | |
| Symmetry imposed | *C*1 | *C1* |
| No. initial particles | 8,352,733 | 6,021,153 |
| No. final particles | 517,853 | 134,422 |
| Map resolution (Å) | 2.32 | 3.04 |
| FSC threshold | 0.143 | 0.143 |
| B factor (Å) | 85.4 | 88.4 |
| **Model refinement** | | |
| Composition | | |
| Atoms | 3,393 | 2,223 |
| Residues | 368 | 268 |
| Water | 511 | |
| Ligands | 1 | 1 |
| Bonds (RMSD) | | |
| Length (Å) | 0.004 | 0.004 |
| Angle (°) | 0.580 | 0.766 |
| Mean B-factors | | |
| Protein | 28.90 | 64.36 |
| Water | 35.69 | |
| Ligand | 28.49 | 45.59 |
| Ramachandran plot (%) | | |
| Favored | 97.27 | 90.23 |
| Allowed | 2.73 | 9.77 |
| Outliers | 0.00 | 0.00 |
| Rotamer outliers (%) | 1.35 | 2.49 |
| Cβ outliers (%) | 0.00 | 0.00 |
| Clash score | 12.73 | 17.85 |
| MolProbity score | 1.85 | 2.59 |
| **Model vs. Data** | | |
| CC (mask) | 0.82 | 0.62 |
| CC (box) | 0.67 | 0.51 |
| CC (peaks) | 0.68 | 0.43 |
| CC (volume) | 0.79 | 0.60 |
| Mean CC for ligands | 0.62 | 0.71 |
